# The mammalian muscle spindle as a tunable feedback controller in locomotion

**DOI:** 10.64898/2026.07.03.736206

**Authors:** Surabhi Simha, Greg Sawicki, Tim Cope, Lena Ting

**Affiliations:** Wallace H. Coulter Department of Biomedical Engineering, Emory University and Georgia Institute of Technology, Atlanta, GA 30332; George W. Woodruff School of Mechanical Engineering, Georgia Institute of Technology, Atlanta, GA 30332; School of Biological Sciences, Georgia Institute of Technology, Atlanta, GA 30332; Department of Rehabilitation Medicine, Division of Physical Therapy, Emory University School of Medicine, Atlanta, GA 30322

**Author notes:** The authors declare no competing interest.

## Abstract

Although muscle spindle sensory signals have been extensively studied, little is known about how and why muscle spindle firing is modulated by the central nervous system during movement. Specialized motor neurons to the muscle spindle, i.e. gamma motor neurons, can profoundly alter spindle firing during behavior, but technological limitations hinder our ability to record gamma motor and muscle spindle sensory signals during most behaviors. We used a biophysical model of a muscle spindle within a muscle-tendon unit to simulate how gamma drive may modulate muscle spindle Ia firing during locomotion. Based on a few available recordings from decerebrate animals, we demonstrate that our model, tuned to passive stretch conditions, can reproduce profound changes in muscle spindle firing in response to identical joint motions in locomotor vs. relaxed stretch conditions. Our model can discover phasic patterns of two types of gamma motor neuron drive based on recorded muscle spindle Ia firing and joint motion. By simulating perturbations, we conclude that: 1) sinusoidal activation of static gamma motor neurons during locomotion, encoding intended movement, modulates muscle spindle signals such that they act as sensorimotor feedback signals based on errors from the intended muscle fascicle length; 2) phasic on/off activation of dynamic gamma motor neurons during locomotion acts as an event detector, heightening muscle spindle Ia responses to discrete perturbations. As such, their muscle-within-muscle structure allows the muscle spindle to act as a highly tunable physical internal model of muscle state to guide movement. Our model supports proposed but as-yet-untested theories of muscle spindle function and offers a framework for extending the testing of muscle spindle function to active, behavioral conditions.

## Introduction

### Muscle spindles are biological sensors for movement control that are modified muscles within muscles, but little is known about the nature of the gamma motor signals that tune sensory signals from muscle spindles during active movements

The common understanding of muscle spindle sensory firing is that it increases as a function of increasing length and velocity (stretch) of its parent muscle-tendon unit, which can be deduced from joint angles (1–4). Much of this understanding comes from experimental conditions where the muscle is relaxed and motion is externally imposed by the experimenter (2). However, muscle spindle firing depends not only on muscle-tendon stretch but also contraction of the muscle (2, 4) and its interactions with the tendon (5–7). Furthermore, muscle spindles are embedded within the muscle fascicles of the parent, i.e. extrafusal muscle, and themselves consist of specialized muscle fibers called intrafusal muscle fibers (1, 2, 8). When motor commands from alpha motor neurons activate and contract the extrafusal muscle, intrafusal muscle fibers may fall slack, reducing or silencing muscle spindle firing and resulting in loss of sensory information during muscle contraction (2, 9). Concurrent motor commands to gamma motor neurons activate intrafusal muscles and can maintain muscle spindle sensitivity during muscle contractions, a phenomenon referred to as alpha-gamma coactivation (1, 2, 4, 9). However, the mammalian muscle spindle is innervated by two independent types of gamma motor neurons—gamma static and gamma dynamic—that have differential effects on muscle spindle firing as measured through experimental stimulation of gamma axons when muscles are stretched in the absence of alpha motor commands (4, 10–12). Furthermore, detailed experiments in reduced animal preparations have found that alpha and gamma motor signals are not always concurrent (13). We currently have little understanding of how gamma motor signals are patterned during active movement, its influence on muscle spindle firing, and effect on movement control.

### Our only empirical evidence of time-modulated gamma motor signals during motor behavior comes from rare recordings during cat locomotion, but the role of these gamma motor signals in generating useful muscle spindle sensory signals for movement control remain untested

Several recordings from locomoting decerebrate or decorticate cats show that gamma static and gamma dynamic motor signals are independently modulated, and thus their activity is not fully explained by alpha-gamma coactivation (13–18). Rare simultaneous recordings of gamma motor activity, alpha motor activity, joint angle, and muscle spindle firing show that relationships between muscle spindle firing and joint angle can vary profoundly between active and passive movement for identical ankle joint movement (Figure 1C). Taylor et al. 2006 (18) demonstrated that gamma static activity is modulated phasically during locomotion, sometimes in advance of alpha motor activity (Figure 1C, row 3). They proposed that gamma static encodes the intended muscle movement, allowing muscle spindle firing to maintain its sensitivity throughout the gait cycle (13, 19, 20). In contrast, gamma dynamic activity is either tonically activated or silent during the gait cycle (Figure 1C, row 4) (13–17) and theorized to signal the timing of extrafusal muscle lengthening, such as in response to contact with the environment, and may increase the sensitivity of the muscle spindle sensory response to such contact (17, 21). Despite these theories, it is currently infeasible to empirically test how gamma motor signals modulate muscle spindle sensory signals to support robust movement control.

**Figure 1:**
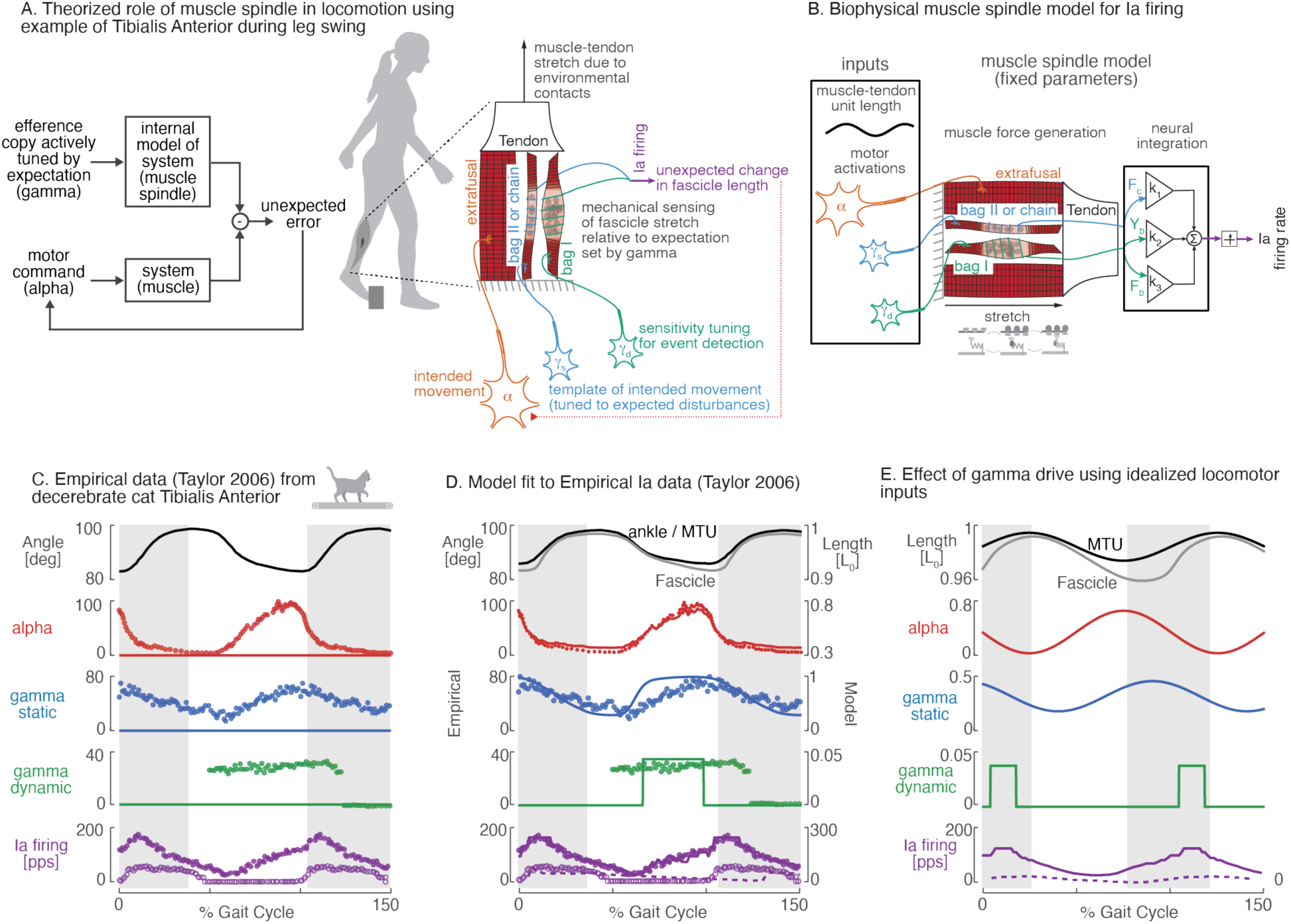
**A.** Block diagram illustrating the movement control loop: motor commands for movement (alpha) drive the muscle system while an efference copy, actively tuned by expectation (gamma), feeds an internal model of the system (muscle spindle). Discrepancies between predicted and actual sensory feedback generate an unexpected error signal. On right is an example during walking: muscle-tendon stretch arising from environmental contacts produces changes in fascicle length. The muscle spindle mechanically senses fascicle stretch relative to expectations tuned by gamma static motor activity, and Ia firing reflects unexpected perturbations. Sensitivity tuning for event detection is set by gamma dynamic activity. **B.** Muscle spindle model consists of a tendon, and parent muscle (extrafusal) and spindle-specific (intrafusal) fibers modelled based on cross-bridge mechanisms, and a phenomenological receptor potential model based on intrafusal fiber force and yank (rate of change of force over time). Model receives alpha, gamma static, and gamma dynamic motor activities, as well as muscle-tendon length as input. All model parameters are kept constant across all simulations. Only inputs are modulated. **C.** Ankle angle (row 1, black), alpha (row 2, red), gamma static (row 3, blue), and gamma dynamic activity (row 4, green) and the corresponding Ia afferent firing (row 5 purple) measured from a cat Tibialis Anterior muscle during decerebrate treadmill walking (bottom row, solid circles) as well as when the same ankle angle was imposed after the cat was anaesthetized (lighter lines for motor activations, open circles for Ia firing) as measured by Taylor et al. 2006. **D.** Model discovers the profile of gamma motor activities when provided with recorded ankle angle, converted to muscle-tendon length, alpha motor activity, and muscle spindle Ia firing (rows 3,5, model lines vs. empirical dots). The profound differences in muscles spindle Ia firing during locomotion (bottom row, solid purple circles) versus when same movement is imposed under anesthesia (bottom row, open purple circles) is predicted by the model by purely changing gamma motor activity (bottom row, solid purple vs. dashed purple line). **E.** Idealized sinusoidal trajectories as inputs to the model to simulate locomotor behaviors discover similar effect of gamma motor activity on muscle spindle Ia firing while allowing for easy parameterization of motor profiles.

### Recently, we developed a biophysical model of a muscle spindle within a muscle tendon unit where the intrafusal fibers receive two types of time-varying and independently modulated gamma motor signals, allowing us to use simulation to test the role of gamma motor modulation in generating useful muscle spindle signals for movement control

The first and fastest feedback from the muscle spindle to affect movement control comes from the spinal stretch reflex—a single synapse connection from the muscle spindle afferent onto the alpha motor neuron (1). For example, when a perturbation stretches the extrafusal muscle that houses the muscle spindle, there is an increase in muscle spindle Ia firing that feeds directly onto the alpha motor neuron of the extrafusal muscle, contracting the muscle, and countering the stretch from the perturbation. Many models have studied how the stretch reflex affects movement control, but very few of them use a muscle spindle model to simulate the stretch reflex. Rather, the models typically assume that the sensory signal is a simple function of either muscle or muscle-tendon length and velocity or joint angle and velocity, inherently eliminating the ability of the simulations to study the role of gamma motor modulation and highlighting the disconnect between the muscle spindle and movement control fields (22–24). Most movement control simulations that do use a muscle spindle model to generate the stretch reflex have relied on phenomenological models of muscle spindles and restricted themselves to ramp-and-hold style, non-cyclic movements, due to the data sets that were used to tune the models (25–28). To simulate a mechanistic muscle spindle Ia sensory signal that can generalize to cyclic movements like locomotion we model the biophysics of two types of intrafusal muscle fibers (bag 1 and bag 2/chain) that receive gamma dynamic and gamma static motor signals, respectively (29, 30). This muscle spindle model is simulated in parallel with a muscle-tendon unit to allow the changing dynamics of the parent extrafusal muscle and tendon to affect the muscle spindle sensory signal, as occurs during active muscle contraction. Notably, the addition of a tendon decouples muscle fascicle and muscle-tendon unit (and joint angle) changes that may also influence muscle spindle firing during muscle contraction. For example, varying alpha and gamma motor signals can explain profoundly different muscle spindle primary signals (Ia afferent firing patterns) observed during isometric contractions based on the relative amplitudes of alpha and gamma motor signals (29). Recent modifications to our muscle spindle model now allow us to extend such simulations to active muscle contractions occurring during cyclic movements to test hypotheses about the role of gamma motor modulation during locomotion (30).

### To evaluate the hypotheses that gamma static modulation serves as a template of intended movement and that gamma dynamic modulation encodes the timing of external contact, we simulated a locomotion-like movement with time-varying gamma motor signals using our biophysical muscle spindle model

We used two methods to determine our model’s ability to reproduce the profound differences in muscle spindle primary signals (Ia afferent firing) due to the effects of gamma motor modulation alone. First, an optimization to empirical muscle spindle Ia firing determined that our model could identify the underlying phasic gamma static and pulsatile gamma dynamic motor signals that generate a given muscle spindle Ia firing (Figure 1D). We then proceeded to use idealized sinusoids to simulate similar profound changes in spindle Ia firing observed in passive stretch versus locomotor conditions for the same change in joint angle and muscle-tendon length (Figure 1E). We show that muscle spindle Ia firing reflects the muscle fascicle movement (Figure 2A) that can be decoupled from muscle-tendon movement (Figure 2B), and that this sensory information can be actively tuned using gamma static motor activity (Figure 2C-E). To test the hypotheses about the role of gamma motor modulation in movement control, we simulated the stretch reflex by feeding the muscle spindle signal at a given time-step as a weighted input to the alpha motor signal to the parent extrafusal muscle on the next time step (Figure 3A). Using such muscle spindle feedback to alpha motor activity, we show that the same total alpha motor activity and muscle fascicle movement can be generated using different ratios of feedforward and feedback alpha motor activity (Figure 3B-D) and that this can be disrupted by modulating gamma static motor signal to deviate from feedforward alpha motor activity (Figure 3E). To more directly test the effect of gamma static motor signals in controlling muscle movement when it deviates from intended movement, we simulated a downhill step (ramp) and a trip (pulse) perturbation during a locomotion-like sinusoidal movement (Figure 4A-B). We found that having a gamma static signal that is a template of the intended movement corrects muscle fascicle length when movement is perturbed away from intended (Figure 4), in a manner consistent with a proportional-derivative control signal (Figure 4). In contrast, the ON-OFF pattern of the gamma dynamic activity as measured during locomotion may signal environmental contact during muscle contraction (Figure 5).

**Figure 2:**
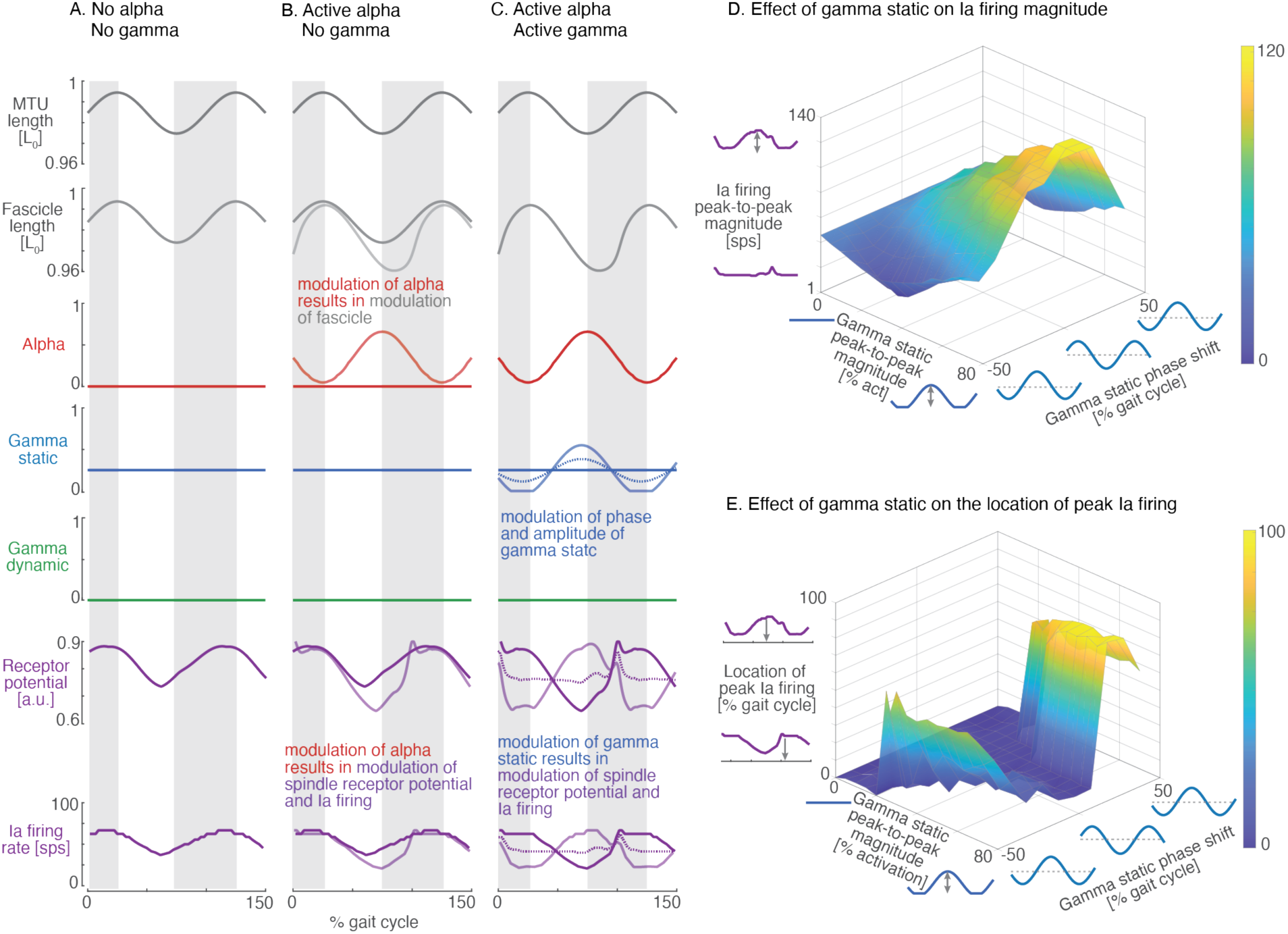
Three simulated conditions are used to show the effect of gamma static motor activity on muscle spindle Ia afferent firing. **A.** Both alpha and gamma static motor activity (row 3 red, row 4 blue) are unmodulated simulating an anaesthetized or relaxed condition where muscle-tendon movement is entirely imposed. Ia firing (bottom row, purple) resembles the shape of muscle fascicle movement (row 2). **B.** Alpha motor activity input is modulated sinusoidally out of phase with muscle-tendon movement to resemble motor activity during locomotion (row 3, light red; dark red from panel A re-drawn for visual comparison). Active contraction due to modulated alpha motor activity causes muscle fascicle movement (row 2, light grey) to deviate from muscle-tendon movement (row 1) and from muscle fascicle movement that was entirely imposed (row 2, dark grey re-drawn from panel A). The change in muscle fascicle movement results in noticeable change in Ia firing (bottom row, light purple vs. dark purple redrawn from panel A). **C.** When gamma static motor activity is also modulated (row 4, blue), in-phase with alpha (row 3, red), Ia firing is reversed (bottom row, light purple) relative to when gamma static activity was absent but muscle-tendon length, muscle fascicle length, and alpha motor activity were identical (bottom row, dark purple redrawn from panel B for visual comparison). An additional condition of differently modulated gamma static activity, keeping all other inputs the same, in dotted blue lines shows that Ia firing profile can also be flattened (bottom row, dotted purple). **D, E.** A heat map shows the effects of varying gamma static phase (x-axis) and magnitude (y-axis) on muscle spindle Ia firing peak-to-peak magnitude in D and on muscle spindle Ia firing phase in E. Gamma static phase was ranged from lagging the sinusoidal muscle-tendon cycle by 50% to leading it by 50% (x-axis range). Note that the muscle-tendon peak occurs at 26 % of the stretch-shorten cycle. Gamma static magnitude ranged from tonically active at 20% to having a peak activation of 70 % (y-axis). Ia firing peak-to-peak magnitude was estimated as the difference between the maximum and minimum firing in a cycle. Ia firing peak location was estimated as the percent of the sinusoidal muscle-tendon cycle (shown as the x-axis in panels A-C) where the peak Ia firing occurred. The Ia firing was passed through a median filter of 5 sample time points to estimate the peak of the cycle rather than short bursts. All inputs apart from gamma static phase and amplitude are fixed across values in D and E.

**Figure 3:**
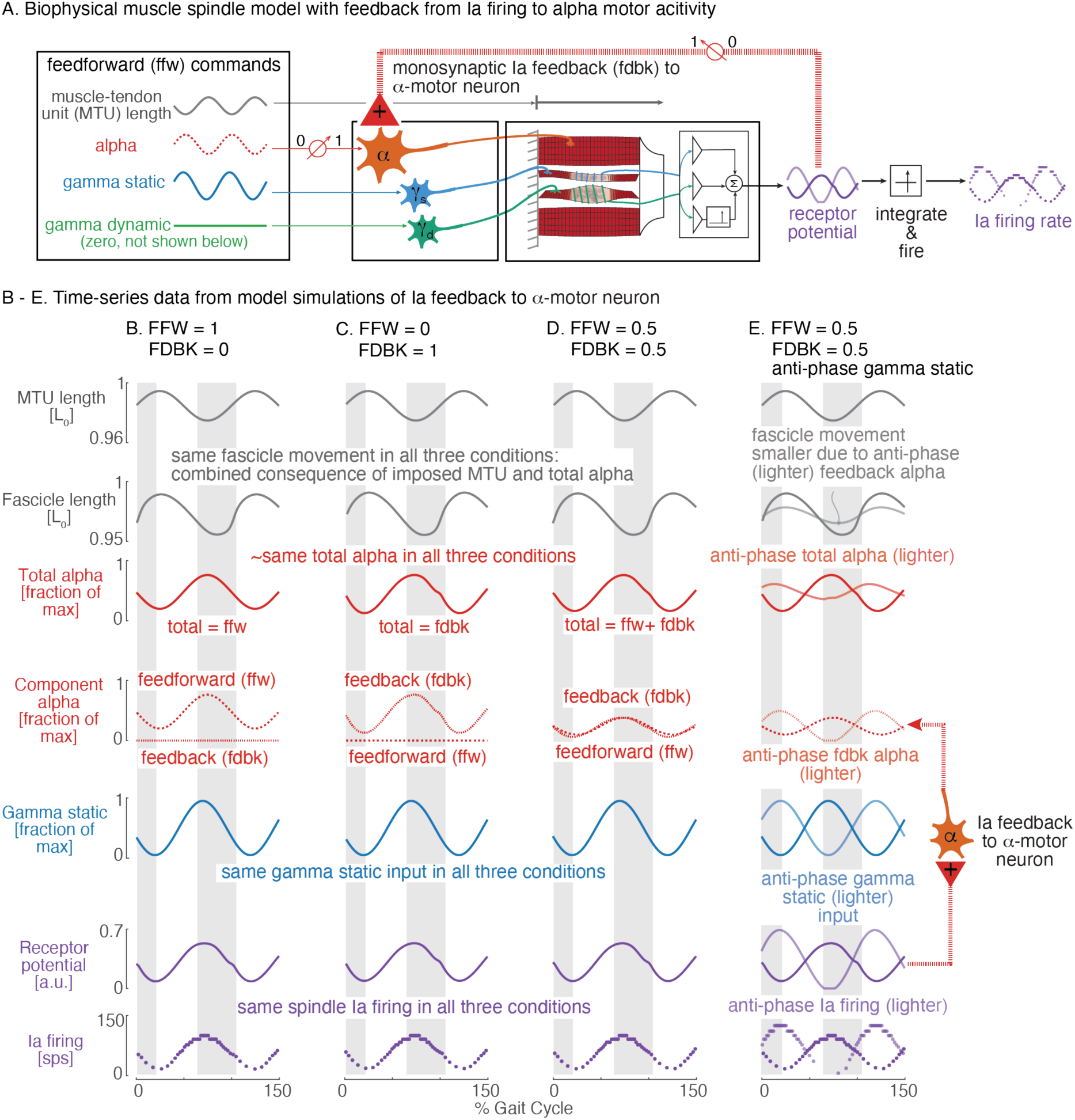
**A.** Schematic of the muscle spindle model with feedback. Feedforward (ffw) and feedback (fdbk) commands drive alpha motor activity, which determines muscle-tendon unit (MTU) and fascicle length. Gamma static motor activity modulates muscle spindle receptor potential, and consequently, Ia firing. Muscle spindle receptor potential is fed back to the alpha motor neuron via the spinal monosynaptic circuit. **B–D.** Simulated locomotor-like sinusoidal cycles across four conditions varying the feedforward-to-feedback ratio (ffw) and feedback gain (fdbk), with gamma static held fixed across conditions. Three different combinations of feedforward and feedback alpha activity (row 4) all result in similar total alpha activity (row 3), fascicle movement (row 2), and muscle spindle firing (bottom row). **E**. Gamma static is phase shifted to be anti-phase (lighter blue, row 5) from the previous conditions (dark blue redrawn from D for visual comparison), causing muscle spindle firing (bottom row, lighter purple; dark purple redrawn from 3D for visual comparison), feedback alpha (row 4, dotted red), and total alpha (row 3, lighter red) to become anti-phase to the previous conditions (darker lines). This results in a small muscle fascicle movement (row 2, lighter grey; dark grey redrawn from 3D) due to monosynaptic Ia feedback. From top to bottom: muscle-tendon length (grey), fascicle length (grey), total alpha (red), feedforward (red solid) and feedback (red dashed) alpha components, gamma static (blue), and Ia firing rate (purple), all plotted as a function of percent sinusoidal cycle of muscle-tendon length.

**Figure 4:**
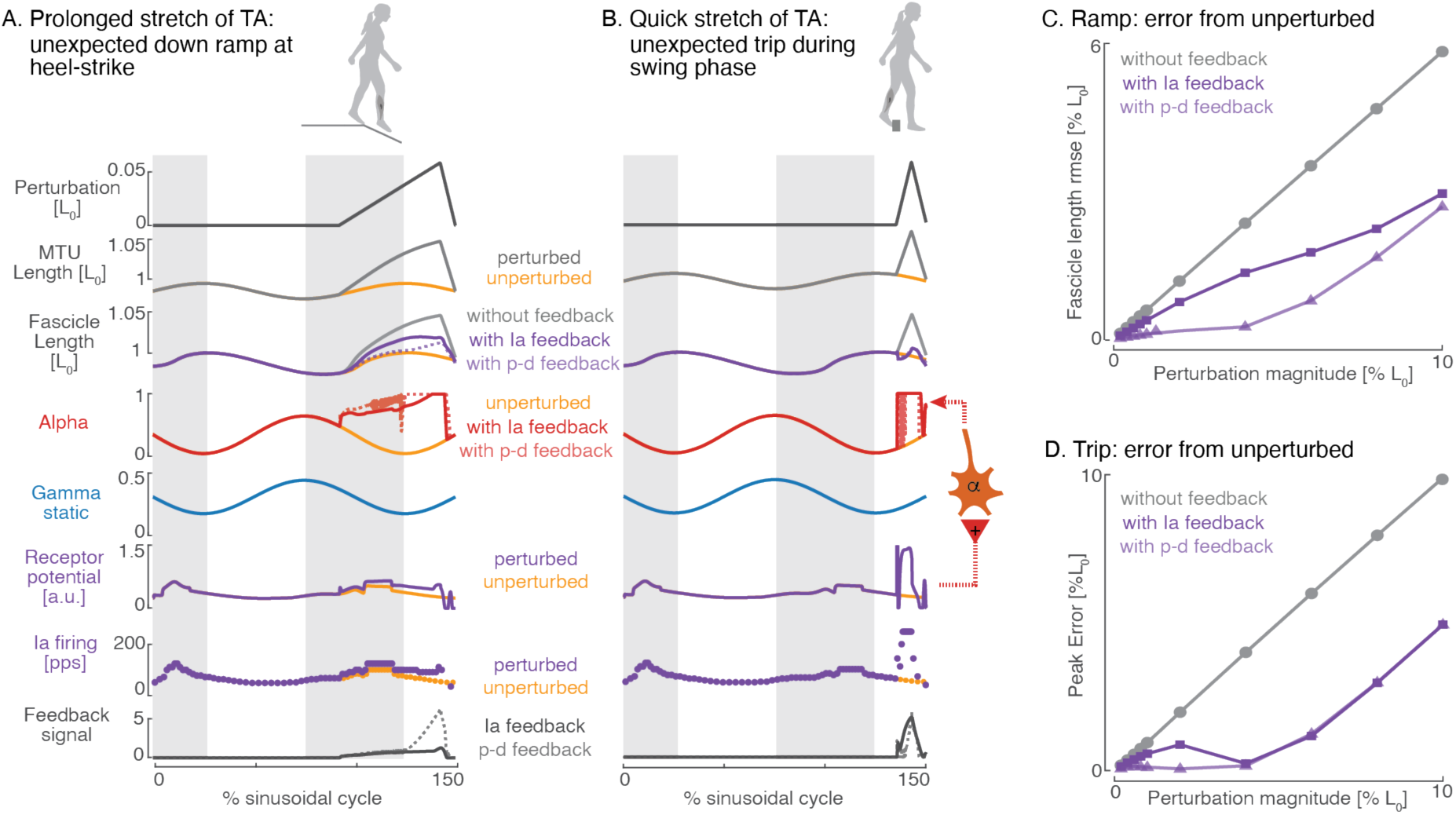
Muscle spindle Ia feedback partially corrects fascicle length deviations in response to muscle-tendon unit perturbations. (A,B). Simulated response to sinusoidal MTU perturbations showing, from top to bottom: perturbation magnitude, MTU length, fascicle length, alpha motor activity, gamma static, receptor potential, Ia firing rate, and feedback control signal, for unperturbed (orange) and perturbed (purple) conditions. Lighter, dashed lines illustrate the effect of a proportional-derivative feedback control on fascicle length. Comparison of fascicle length (row 3) with (purple) versus without (grey) feedback control across perturbation types, shows that Ia feedback attenuates but does not eliminate fascicle length deviations from unperturbed (orange). (C,D) Peak fascicle error (for trip-like, pulse perturbation) and fascicle length RMSE (for downhill, ramp) as a function of perturbation magnitude, with (purple) and without feedback (grey) control, shows that feedback reduces but does not eliminate length deviations across a range of magnitudes. Ia feedback driven fascicle length regulation is similar to that from a proportional-derivative feedback controller when the perturbation is rapid, trip-like. For slower ramp-like perturbations, Ia feedback reduces fascicle deviation but underperforms relative to the proportional-derivative feedback controller at smaller perturbation magnitudes.

**Figure 5:**
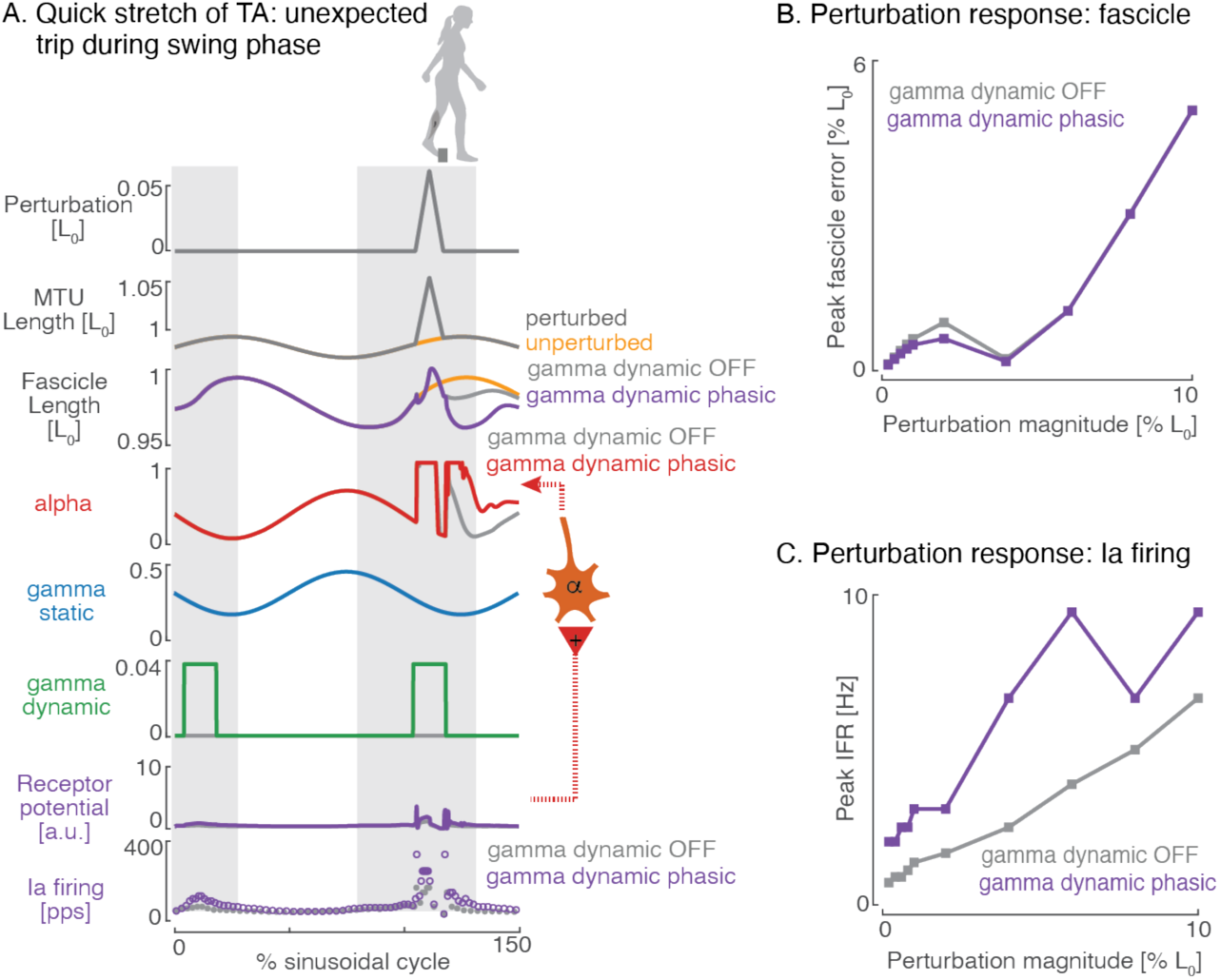
Gamma dynamic motor activity modulates Ia firing sensitivity to MTU perturbations but has limited effect on fascicle length correction. (A) Locomotor cycles with MTU perturbations were simulated with two conditions of gamma dynamic activity: absent gamma dynamic (grey lines) and when gamma dynamic was active for part of the gait cycle as determined in Figure 1E (row 3 purple, row 4 red, row 6 green, row 7,8 purple). From top to bottom: perturbation magnitude, MTU length, fascicle length, alpha motor activity, gamma static, gamma dynamic, receptor potential, and Ia firing as a function of percent gait cycle. Gamma static profiles are matched between conditions, isolating the effect of gamma dynamic modulation. Having gamma dynamic active consistently increased the muscle spindle Ia firing to perturbations (A, bottom row, purple circles higher; C, purple greater than grey). However, this did not translate to any further reduction in the deviation in fascicle length (A, row 2, B). B-C. These trends remained at a range of perturbation magnitudes.

## Results

### Fitting simulations discover locomotor gamma motor activity

#### To simulate the interactions between alpha and gamma motor activity on muscle spindle Ia firing during movement, we imposed periodic length changes to a biophysical muscle-tendon model

The model received muscle-tendon length and motor activations as inputs and estimated the length and force of the extrafusal muscle fascicle using differential equations to generate the cross-bridge attachment and detachment rates according to a Huxley-type muscle model (29–31). The extrafusal muscle fascicle length was then applied to the two spindle-specific intrafusal muscle fibers (one bag 1 and one bag 2/chain) to estimate the force from each fiber (Figure 1B). The intrafusal muscle fibers were also modelled according to a Huxley-type muscle model but differ from the extrafusal muscle and each other in their parameters (29, 30, 32, 33). The forces from both intrafusal fibers and the rate of change of force, termed yank, from the bag 1 fiber were combined in a non-linear transformation to estimate the receptor potential at the spike initiating region. Finally, the receptor potential was integrated to determine the muscle spindle Ia afferent firing. While prior published versions of this muscle spindle model only reported the receptor potential, we report afferent firing in all the main results for easier comparison to empirical data (29, 30).

#### Based on muscle spindle Ia firing recorded during locomotion, our optimization discovered time-varying gamma static and gamma dynamic motor patterns that closely resemble the recorded motor signals

In accordance with other findings (14, 15, 17), Taylor et al. 2006 found that gamma static and gamma dynamic motor activity were differently modulated during locomotion in decerebrated cats (Fig. 1C). By using an optimization algorithm that 1) took in gamma static and dynamic motor activity as tunable parameters, 2) took in the recorded alpha motor activity and muscle-tendon length (derived from recorded ankle angle) as fixed input, and 3) was required to fit the model output to the recorded Ia firing, we found that our model identified the locomotor gamma motor activity that fit the empirical Ia firing with a Pearson’s r = 0.97 (Fig. 1D). Specifically, gamma static signal discovered by the optimization was phasic and slightly in advance of the alpha motor signal, qualitatively resembling empirical data (Pearson’s r = 0.75). The qualitatively similar shape between the model and recorded data is notable because gamma static was parameterized as a 5-point spline with a variable phase, which meant that search space for the optimization included constant tonic activity, absent activity, or any shape that a 5-point spline can generate. In contrast, the gamma dynamic signal was parameterized by two variables determining the onset and offset time of gamma dynamic activity, and one variable determining the magnitude of activation. The gamma dynamic timing discovered by the optimization overlapped with that of the recorded signal but with a slightly delayed onset and early offset (Pearson’s r = 0.38).

#### Extinguishing motor activity for muscle tendon length identical to the locomotor condition dramatically alters the pattern of muscle spindle firing, resembling muscle spindle activity in anaesthetized animals

Taylor et al. 2006 also found that muscle spindle Ia firing showed large qualitative changes when the ankle movement measured during locomotion was imposed on the same cat when it was anaesthetized (Figure 1C, bottom row, open circles). Specifically, the muscle spindle Ia firing increased during muscle shortening in the locomotor condition while it was silent in the anaesthetized condition, and the firing decreased during muscle stretch during the locomotor condition while it remained nearly constant or increased in the anaesthetized condition (Figure 1C, bottom row, closed vs. open circles). We simulated the anaesthetized condition by setting the motor inputs to the model (alpha, gamma static, and gamma dynamic) to zero while applying the same muscle tendon length as the locomotor condition. The muscle spindle Ia firing from our model displayed a similar qualitative difference between the firing from the locomotor condition—in the anaesthetized simulation, firing increased during stretch but was silenced during shortening (Figure 1D, bottom row, dashed lines).

#### Simplifying the model inputs to idealized trajectories generated similar muscle spindle Ia patterns in locomotor and passive conditions, enabling parameter variations for hypothesis testing in predictive simulation

To more conceptually test the role of gamma motor activity to affect the muscle spindle Ia firing, we used an ideal sinusoidal muscle-tendon length as input to the model. We simulated two conditions: anaesthetized and locomotor. In the locomotor condition, we commanded a sinusoidal profile of motor activity based on Taylor 2006 findings: alpha activity was a sinusoid out of phase with the muscle-tendon length and gamma static was slightly phase advanced from alpha by 20 % of the gait cycle. Gamma dynamic timing was offset relative to the empirical measures, and we suspect that this reflects some of the effect of the muscle fascicle kinematics on Ia firing, since the muscle kinematics no longer match the empirical values (Figure 1 D vs. E, row 1). In the anaesthetized condition, alpha and gamma motor activity were maintained at zero throughout the stretch-shorten cycle. The effect of gamma motor activity, as observed by qualitative differences between the locomotor and anaesthetized conditions, was similar to when muscle-tendon length matched empirically recorded data (Figure 1E, bottom row, solid vs. dashed lines). Specifically, the muscle spindle firing increased during muscle shortening in the locomotor condition while it was silent in the anaesthetized condition, and the firing decreased during muscle stretch during the locomotor condition while it remained nearly constant or increased in the anaesthetized condition. Such idealization of the input trajectories allows us to easily parameterize gamma static activity using phase and amplitude to evaluate hypotheses about its role in movement control.

### Gamma static may serve as a template of intended muscle fascicle movement

#### Muscle spindle Ia firing reflects muscle fascicle movement that can be decoupled from that of muscle-tendon unit due to alpha motor activity

Since muscle fascicle is the controllable unit for the nervous system through alpha motor activity, and muscle spindle Ia firing can only control movement by influencing alpha motor activity, we hypothesized that muscle spindle firing would better reflect the muscle fascicle movement compared to muscle-tendon movement. We simulated two conditions, both with identical sinusoidal muscle-tendon length input and zero gamma motor activity (Figure 2 A, B, rows 1,4,5). In the first condition, alpha motor activity was set to zero while the second condition used a sinusoidal alpha motor activity that was out of phase with the muscle tendon movement to simulate a locomotor condition (Figure 2 A, B, row 3). When alpha motor activity was absent, muscle fascicle movement resembled muscle-tendon movement (Figure 2A, row 1 vs 2) but in the presence of alpha motor activity, the muscle fascicle movement deviated (Figure 2B, row 1 vs 2, light red alpha causes light grey muscle fascicle movement). The change in muscle fascicle movement caused due to alpha motor activity resulted in muscle spindle Ia firing to change as well (Figure 2B, bottom row, light purple vs. dark purple redrawn from 2A for visual comparison). Specifically, deviation can be observed in both the muscle spindle Ia firing and muscle fascicle at the onset of stretch, but not in the muscle-tendon unit length (Figure 2B).

#### Modulating gamma static motor activity can reverse or flatten muscle spindle firing for identical alpha activity and muscle-tendon movement

To demonstrate the effect of gamma static activity on muscle spindle Ia firing, we simulated a third condition where gamma static was modulated sinusoidally with alpha activity and out of phase with muscle-tendon length to simulate a locomotor condition (Figure 2C). Gamma dynamic activity was maintained at zero to highlight the role of gamma static activity alone. Phasic gamma static activity was able to reverse the muscle spindle Ia firing, relative to when gamma static was unmodulated (Figure 2C bottom row, light solid lines. Dark solid lines re-drawn from B for visual comparison). Gamma static activity modulation could also reduce muscle spindle Ia firing to flatten out the modulation (Figure 2C, bottom row, dashed). A sweep across a range of gamma static phase and amplitude found that muscle spindle Ia firing can be profoundly modulated by gamma static motor activity to the muscle spindle even when all other inputs are identical (Figure 2D, E). Such modulation of muscle spindle Ia firing by gamma static activity when all other inputs to the muscle spindle are unchanged suggests that the muscle spindle is not a passive sensor of muscle fascicle movement but rather contains information about the muscle fascicle state, that can be tuned by gamma static motor activity.

#### The muscle spindle Ia firing can modulate muscle fascicle movement through the stretch reflex driven feedback to alpha motor activity

To simulate the monosynaptic stretch reflex, we converted the muscle spindle Ia receptor potential from our model at each time step into a scaled, non-negative feedback signal that was added to the alpha motor activity on the next time step (Figure 3A). We then evaluated the ability of the muscle spindle Ia feedback to modulate the muscle fascicle length by using a sinusoidal muscle-tendon length as input to simulate a locomotion-like condition. Alpha motor activity was input as a sinusoid out of phase with the muscle-tendon length and this was now defined as the *feedforward* alpha motor activity (Figure 3A). The total alpha motor activity to the model now also consisted of a *feedback* component that was the scaled, non-negative muscle spindle Ia output (Figure 3A). Gamma static motor activity input was a signal in phase with the feedforward alpha motor activity while gamma dynamic activity was maintained at zero to focus on the effect of gamma static activity (Figure 3B-D, row 4 blue). Only in these set of simulations, we introduced two additional parameters: feedforward gain and feedback gain, to vary the relative ratios of the feedforward and feedback alpha motor activity to the total alpha motor activity. By setting the feedback gain to zero, we first report the muscle fascicle movement when the total alpha motor activity is entirely feedforward (Figure 3B). Next, by setting the feedforward gain to zero, we show that the same total alpha motor activity and muscle fascicle movement can be obtained purely from muscle spindle Ia afferent feedback (Figure 3C). Finally, we simulate a condition most likely to match reality where both feedforward and feedback alpha motor activity contribute to the total alpha motor activity, such that the total alpha motor activity and muscle fascicle movement match the earlier conditions. These set of results highlight the range of effects that muscle spindle Ia firing can have in modulating muscle fascicle movement—from not at all to replacing the effect of feedforward activity or anything in-between.

#### Gamma static motor activity can modulate muscle fascicle movement through its effect on the muscle spindle Ia firing that generates the feedback alpha motor activity

To evaluate the effect of gamma static motor activity on modulating muscle fascicle movement, we shifted the phase of gamma static activity such that it was now out of phase with the feedforward alpha motor activity (Figure 3E, row 4 blue). The total alpha motor activity was generated using the same ratio of feedforward and feedback alpha motor activity as when gamma static activity was in phase with alpha motor activity (Figure 3D). The resulting total alpha motor activity was now out of phase with the feedforward alpha motor activity (Figure 3E, row 3, light red; dark red redrawn from 3D for visual comparison). This means that the total alpha activity is now increasing and generating a muscle fascicle contraction when the imposed muscle-tendon movement is lengthening. Such opposition from the total alpha motor activity to the movement imposed by the muscle tendon on the muscle fascicle results in the muscle fascicle moving less relative to all the prior conditions when total alpha motor activity aided the imposed movement (Figure 3E, row 2, light gray; dark grey redrawn from 3D for visual comparison).

#### When gamma static activity acts as a template for intended movement, deviations in muscle fascicle length from intended movement are reduced

To evaluate the effect of the muscle spindle-mediated stretch reflex in maintaining intended muscle fascicle movement, we simulated two types of deviations to the muscle fascicle movement (perturbations): unexpectedly stepping onto a downward sloped terrain and tripping. The perturbations were simulated by applying stretches to the muscle-tendon movement. Downhill was simulated with a slow, constant-velocity ramp stretch (Figure 4A) acting on the Tibialis Anterior around heel-contact when stretch is expected, and tripping was simulated as a rapid, brief stretch acting on the Tibialis Anterior during mid-swing (Figure 4B). We used the alpha motor activity and gamma static motor activity from the idealized-trajectory locomotor simulations (Figure 1E) and treated the resulting muscle spindle Ia receptor potential as the baseline, steady-state Ia output. Deviations from the baseline muscle spindle Ia receptor potential were used as low-pass filtered, scaled, non-negative feedback to the alpha motor activity. The additional muscle spindle feedback driven alpha motor activity (Figure 4A,B, row 4, dark, solid red) reduced the muscle fascicle length deviation away from the intended steady-state movement (Figure 4A,B, row 3, 4; C,D, dark, solid purple), where the intended movement is assumed to be the unperturbed movement (Figure 4A,B, row 3, orange). We further found that when gamma static activity acts as a template of intended movement, the effect of muscle spindle Ia firing on muscle fascicle length is comparable to a proportional-derivative feedback control signal that is acting to maintain the intended muscle fascicle length (Figure 4A,B, rows 3 lighter, dashed lines; Fig 4C,D, light purple). A sweep across a range of perturbation magnitude found a similar effect of Ia feedback reducing muscle fascicle deviations (Figure 4 C,D).

### Gamma dynamic may help in signaling of limb contact with environment

#### Phasic gamma dynamic activity results in larger muscle spindle Ia firing in response to discrete perturbations

Since gamma dynamic motor activity during locomotion is pulsatile, being active only during muscle contraction, we simulated two conditions to test the effect of locomotor gamma dynamic activity timing on muscle spindle Ia firing: one where gamma dynamic was always absent and one where it was active during only part of the gait cycle as discovered by the optimization (ref. figure 1E), and as observed during locomotion. Gamma static activity, alpha activity, and muscle tendon movement were fixed to match the values identified in the locomotor condition (Figure 1E). Since gamma dynamic activity is known to modulate response to rapid stretch, we focused on the brief “pulse” stretches applied to the muscle tendon. The presence of gamma dynamic activity noticeably increased the muscle spindle Ia firing in response to perturbations relative to when gamma dynamic activity was absent (Figure 5A bottom row, 5C; purple vs. grey). Shifting the perturbation as well as gamma dynamic activity to different phases of the gait cycle produced a similar result.

#### Gamma dynamic mediated muscle spindle Ia firing to perturbations only marginally contributed to muscle fascicle error correction

To directly test the effect of locomotor gamma dynamic activity in correcting muscle fascicle error through the stretch reflex, we evaluated the role of the three gamma dynamic timing conditions in affecting muscle fascicle length following perturbations. Interestingly, the corrective change in muscle fascicle length due to gamma dynamic activity, in the presence and absence of gamma dynamic activity did not differ (Figure 5A row 3, 5B; purple vs. grey). One explanation for this observation is that our filtered feedback of Ia receptor potential to generate a reflex does not appropriately capture the dynamics of the process of converting spikes to muscle force—rapid doublet spiking can result in enhanced force from the activated muscle (34, 35).

## Discussion

### Taken together, our simulation results support as-yet-untested theories that gamma static and gamma dynamic motor activity during locomotion tune muscle spindle Ia signal to act as both a feedback controller and an event detector

Supporting existing hypotheses that two, independently modulated, gamma motor signals shape muscle spindle Ia firing, our biophysical muscle spindle model discovered the distinct sinusoidal modulation of gamma static versus on-off modulation of gamma dynamic signals based on recorded empirical data from cat locomotion (13, 16–18, 36–38). The parameters of our muscle spindle model were only tuned to muscle spindle Ia firing measured from anaesthetized cats (17, 39, 40), when movement was externally imposed on the muscle-tendon unit. Consequently, our modelling results highlight the profound differences in muscle spindle firing that arise purely from changes in gamma motor activity, in accordance with empirical recordings (4, 10, 11, 40–43). Our biophysical formulation of the muscle spindle in parallel with muscle-tendon unit enabled the model to generalize to active, behavioural conditions, and further to in silico testing of theories about muscle spindle function during locomotion. Our findings support the idea that gamma static motor activity can shape the degree of feedback-mediated alpha motor activity to the extrafusal muscle (1, 4, 44, 45) during locomotion (46), while also tuning a feedback controller to regulate muscle fascicle length based on the intended movement (13, 20, 23). In contrast, gamma dynamic activity may tune the sensitivity of the rapid burst of muscle spindle firing (10, 17), signaling contact with the environment (17, 21), that require rapid muscle force generation (34, 35) or changes in motor drive to the muscle (47, 48). Our results here support the theories that muscle spindles can act as task-dependent, tunable, peripheral internal models with independent alpha-gamma activation (4, 49, 50), enabling effective movement control in familiar (21) and novel environments (51, 52), and highlight the need for empirical testing of muscle spindle function during awake behaviour.

### Despite sophisticated theories that gamma drive tunes muscle spindle function, whether it does so during intact movement remains unresolved—a gap that simulations have recently begun to address

Extensive neurophysiology experiments in animals have directly measured gamma motor activity during awake behaviour to find that it is modulated independently of alpha motor activity (13–18), and have inferred that gamma activity modulation is task-dependent (50). However, it is only recently that muscle fascicle kinematics have begun to be recorded simultaneously with muscle spindle firing (7), and not yet in awake movement. Since muscle fascicle kinematics are a primary factor affecting muscle spindle firing (4, 53), and they can be decoupled from joint angle during movement (5, 54, 55), the effect of gamma motor activity on muscle spindle firing cannot be known without also knowing muscle fascicle kinematics. Furthermore, these experiments are highly invasive, and therefore cannot be performed in intact, healthy animals (13, 43). A separate line of work in healthy, awake humans has suggested the presence of gamma motor drive that is independent of alpha (56, 57), albeit indirectly inferred through measurements of muscle spindle firing (58)—except in one case (56, 59)—since gamma motor neurons are not easily accessible in minimally invasive experiments (9). Notably, these experiments have largely found minimal effect of independent gamma motor drive on muscle spindle firing (58, 60, 61). However, the experiments in humans require the use of fine microelectrodes to remain lodged within afferent axons of ∼10 μm diameter and thus are highly sensitive to large or voluntary movements (62). Finally, similar to animal studies, only one study recorded muscle fascicle kinematics concurrently with muscle spindle firing, and only in relaxed, externally imposed movement (6). Given these limitations on empirical data, the debate on the relevance of independent gamma drive on movement control remains unresolved (9, 63–65). As we show through the following discussion of our results and existing literature, independent gamma drive during movement is most likely to have observable effects on muscle spindle firing in non-steady-state motor tasks—behaviour that is the least studied thus far. The few studies in human movement that have tested non-steady-state motor tasks have found evidence consistent with the theories of task-dependent modulation of muscle spindle firing using gamma motor drive (51, 52, 57). Technical limitations still prevent in-depth empirical testing, though there is some development through the use of optogenetics in mouse models (66). In the meantime, there has been a surge in muscle spindle modelling efforts recently, which can be leveraged to design targeted experiments that improve our understanding of muscle spindle function in movement control (29, 30, 67–71).

### Our biophysical modeling results highlight that vastly different sensory signals—despite similar measures of biomechanics and muscle activations—can arise from descending modulation of the muscle spindle

A key feature of our muscle spindle in muscle-tendon model that enabled this study is the model’s biophysical formulation that affords generalizability beyond the phenomenon to which it was tuned. In this and prior versions of our biophysical muscle spindle model, muscle spindle responses were tuned only to conditions of quiescent alpha and gamma drive, yet that was sufficient to produce realistic firing in active isometric, lengthening, and shortening conditions (29). Beyond the anatomical arrangement of the muscle spindle, muscle, and tendon, our biophysical formulation includes muscle properties such as actin and myosin molecular properties, mechanotransduction of the sensory signal, and distinct roles of the different spindle-specific fibers. These biophysical properties give rise to classically observed properties of muscle spindles, such as the initial burst, plateau, and decay in firing with stretch, history dependence, power-law relationships of firing to stretch velocity (29, 30, 72). The model’s generalizability lets us show that independent control of gamma-static and alpha drive allows the spindle to encode muscle fascicle kinematics distinct from muscle-tendon unit length—a distinction absent in many prior models that either used a fixed gamma activity (73–76), gamma drive co-activated with alpha (22, 23, 27, 77) or always coupled muscle fascicle and tendon kinematics (22, 23, 76). To our knowledge, only one prior modelling study has inferred independent, time-varying gamma-static and gamma-dynamic drive, doing so by fitting muscle spindle Ia firing during active human movements without access to the underlying gamma motor activity (28). Here, leveraging cat locomotion, where gamma motor activity has been directly recorded (18), we tested the model-predicted gamma-static and gamma-dynamic drive against empirical recordings. Finally, by simulating the monosynaptic reflex pathway, we ran in silico experiments to test the potential roles of gamma static and gamma dynamic activity in determining muscle spindle Ia firing and movement control.

### Gamma static drive may serve to reinforce feedforward motor commands in well-learned tasks and generate sensorimotor error signals to facilitate motor learning for new tasks

Muscle activity during steady-state locomotion consists of both feedforward motor activity driven by Central Pattern Generators (78) as well as motor activity generated based on contributions from sensory feedback that is processed at various levels of the neural circuitry (79–81). Here, we show that when gamma static motor activity is periodically modulated at the locomotor alpha motor frequency, monosynaptic feedback from the muscle spindle Ia afferent can create a biologically plausible feedback component of the total muscle activity (Figure 3 B-D). While monosynaptic feedback from the Ia afferent onto the parent alpha motor neuron is reduced for plantar flexors during walking through presynaptic inhibition (82, 83) such suppression is muscle-and task-dependent (84, 85), permitting for larger contributions from the monosynaptic pathway during tasks like standing (83, 86, 87). We evaluate the validity of the model using empirical data from the cat Tibialis Anterior muscle but our conceptual tests on the role of gamma motor activity during movement, such as in shaping the feedback muscle activity, as task-and-muscle agnostic. Our results highlight the effect of tuning gamma motor activity, which is different for different muscles and tasks. Furthermore, gamma static activity also modulates group II afferent firing from the muscle spindles (4, 40, 42, 88) that is implicated in the modulation of alpha motor activity during walking (89, 90). Thus, when gamma static and alpha are congruent, such as in a well-practiced steady-state locomotion task, the muscle spindle feedback can reinforce the feedforward alpha motor pattern. On the other hand, our simulations show that when gamma-static and alpha motor activity are out of phase, such as when learning a new task, feedback and feedforward alpha motor activity may be discordant, resulting in unintended muscle spindle firing (Figure 3E). Consistent with this, muscle spindle firing during visuomotor learning is modulated independently of muscle activity and kinematics (51, 52), increasing during active adaptation in a manner that predicts learning rate (51) and decreasing as performance improves and proprioceptive feedback realigns with the task (52). Thus, state-dependent muscle spindle feedback could facilitate motor learning where the increased afferent signal during incongruence may flag a persistent sensorimotor error, and re-tuning gamma-static toward the new task, e.g., when adapting from walking on hard ground to sand, restores low firing once the updated feedforward command is congruent with the desired movement. The process of learning the new motor commands may be through local gradient descent along a high-level cost value, identifying an economical pattern (91–94). However, the mechanisms of sensing the gradient are still unknown (95–97) and furthermore, not everyone always appropriately updates their motor commands in response to a change in task (95, 96). Muscle spindle firing may serve as a peripheral signal that helps the central nervous system associate the change in sensory signals with appropriate adaptation of motor commands.

### Gamma static drive may be tuned to regulate muscle fascicle length during locomotion and tuned to regulate other muscle variables according to task requirement

Well-learned gamma static motor patterns during locomotion may be shaped by the need to regulate muscle fascicle length (54, 98, 99) about the intended movement (100), avoiding interpreting sensory information through slower, longer-latency pathways (101–103). Our simulations show that the monosynaptic spindle feedback to alpha motor neuron can help regulate muscle fascicle length (Figure 4). As muscles typically act in series with elastic tendons, fascicle length can be regulated by maintaining the muscle tension (104), consistent with the muscle spindle receptor potential being well-represented by a mechanotransduction of intrafusal fiber force and its time derivative, yank (29, 30, 72, 105, 106). Thus, muscle spindle sensory signals can be decoupled from the biomechanical state of the joint or muscle-tendon, encoding the decoupling between state of the muscle fascicle and the joint. While our simulations here use idealized sinusoids, plantar flexor muscles often act isometrically during stance in steady state walking (54, 98), suggesting rapid muscle length regulation may be beneficial. Maintaining isometric fascicle length during locomotion can enable the muscle to generate more force per unit of activation due to two reasons: 1) force decreases when a muscle shortens due to the force-velocity relationship (107) and 2) length changes alter the actin-myosin overlap (108), reducing the number of attached cross-bridges (109). However, not all tasks, or even all muscles in the same task, benefit from maintaining isometric muscle fascicle length or even regulating muscle length to generate high force. For example, tasks like cycling (110) or walking uphill or downhill (111, 112) depend on performing net positive or net negative work and may benefit from regulating muscle force or velocity, rather than length. Here, gamma static activity may be calibrated to expected task stiffness such that the muscle spindle firing reports the unexpected external force acting on the muscle. Task-dependent regulation of the balance between monosynaptic (Ia only) and polysynaptic pathways (83, 113, 114) could weight and combine the muscle spindle Ia, II, and the Golgi Tendon Organ Ib signals to generate estimates of deviation from expected muscle mechanical work. Accordingly, group Ib and group II afferents converge onto shared premotor interneurons (115), potentially allowing for correlational estimates of mechanical work (116, 117). We focused on level-ground locomotion here since the only simultaneous recording of gamma motor activity and muscle spindle firing is from locomotion (18). More broadly, precise tuning of the gamma static motor activity according to muscle function and task may allow for rapid, task-relevant feedback signals relative to intended behaviour.

### Gamma dynamic motor activity may be used to tune the muscle spindle sensory signal to serve as a salient and temporally precise signal of critical events such as ground contact during locomotion

Our simulations find that while gamma dynamic activity modulates muscle spindle firing in response to perturbations, this additional activity does not lead to further improvement in feedback regulation of muscle fascicle length beyond that provided by gamma static activity. However, empirical recordings from cat locomotion have consistently found that gamma dynamic activity begins around the onset of alpha motor activation and is silenced shortly after the onset of stretch (14, 15, 17, 18), after ground contact would have occurred, suggesting a specific function of gamma dynamic timing. One hypothesis about the function of gamma dynamic activity during locomotion is that it generates a rapid, salient sensory burst in response critical events involving contact with the physical environment (13, 17, 18, 50). The neurophysiology of gamma dynamic motor neuron supports this hypothesis: it only innervates the viscoelastic bag 1 intrafusal fiber that has a slow creep-response to stretch (4, 41, 118, 119), resulting in the muscle spindle Ia firing being primarily responsive to rapid stretches in the absence of gamma static activity (10–12, 120). Thus, the salient muscle spindle firing sensitized by gamma dynamic activity to rapid perturbations may generate a faster muscle force response (34, 35), or even trigger a change in higher levels of neural control such as brainstem mediated responses as seen in postural control to perturbations (121, 122), or even a cortically mediated response to perturbation during posture or locomotion (101, 123).

### The muscle spindle may thus be a tunable biomechanical internal model that uses physical computation to encode the mechanical consequences of body-environment interactions on the desired movement

Conventionally, central internal models are assumed to estimate the consequences of physical interactions (124) through neural computations, which incur both processing and transmission delays (103). The highly nonlinear dynamics of the body’s interactions with the environment are especially difficult to compute (125), challenging even modern-day models of soft body contacts and collisions that are critical to terrestrial locomotion (126). Further, muscle force itself is highly non-linear and history dependent (127) such that the muscle force cannot be known simply by its level of activation by the nervous system but depends upon both the current and prior muscle state, its activation, and the external forces acting on it (128, 129). The muscle-within-muscle design of the muscle spindle sensory organ acts as a physical model of the extrafusal muscle that directly experiences the muscle’s dynamics resulting from both muscle-generated force and external forces without any delays, serving as a mechanical model of the muscle. Beta motor neurons send collaterals to the muscle spindle’s intrafusal muscle, enabling them to have the same central drive as the parent muscle (130, 131); amphibians, and reptiles only have beta drive (132). Gamma motor neurons and muscle spindle Ia afferents with highly transient firing responses are a feature of mammals and avians, where dynamic, limbed terrestrial locomotion is sensitive to long neural feedback delays (132). As such, gamma motor neurons may receive descending inputs from area 3a of the somatosensory cortex (133), suggesting a pathway between cortical areas interpreting muscle spindle feedback to directly alter their primary sensory output. Similar to theories of motor learning, such a cortical pathway operating in parallel to the spinal pathways and may serve as a teaching signal, particularly during novel motor tasks (134). Consistent with this, spindle dynamic sensitivity is upregulated during difficult or unfamiliar tasks (135). Similar modified effectors operating as sensory organs can be observed in flies, where halteres are a very small, modified wings that operate alongside flight wings, for the control of flight, another highly dynamic and mechanically complex movement (136). Moreover, haltere outputs act monosynaptically on the motorneurons of the wing-steering muscles (137), as well as receive top-down motor inputs such as from visual pathways (138). Broadly, our results support hypotheses that muscle spindles can be tuned to support bottom-up and top-down sensorimotor processes, allowing the muscle spindle to play complex and varied roles during different movements and contexts.

## Methods

All code was written in MATLAB 2022b (Mathworks, PA). Simulations were run at 1000 Hz. Equations are provided in Simha and Ting 2024.

### Muscle spindle model

Our model consists of the muscle-tendon unit (parent, extrafusal muscle fascicle with the tendon in series), two spindle-specific, intrafusal muscle fibers, and the mechanotransduction process. We model both the intrafusal and extrafusal fibers using similar crossbridge mechanisms but use different parameters to simulate the differences in their force generating properties. Length and activation are imposed on the muscle-tendon unit, and the corresponding extrafusal muscle fascicle length change is estimated. This muscle fascicle length is then applied to the intrafusal muscle fibers. Similar to our previous models, we use a phenomenological model of the mechanotransduction process: a weighted sum of the force and rate of change of force (yank) from the intrafusal fibers generates the total receptor potential at the spike initiating region. The receptor potential is integrated over a fixed time window to generate the muscle spindle Ia firing rate.

### Extrafusal muscle model

We modelled the extrafusal muscle as a half sarcomere in series with a tendon. The tendon is modelled as linear spring while the half sarcomere is simulated using differential equations of the cross-bridge dynamics that govern force production in a sarcomere. We use a modified version of the standard Huxley equations that allow the myosin heads on the thick filament to exist in one of three states: attached, detached, super-relaxed (139), as published in Simha and Ting 2024 (30).

### Intrafusal muscle model

The contractile properties of the intrafusal fibers were also modelled based on cross-bridge dynamics and the passive parallel element was modelled as linear springs (29, 30, 140). For the intrafusal fibers we used a simpler muscle model where the myosin heads only exist in one of two states: attached to the actin sites or unattached, as published in Simha and Ting 2024.

### Receptor potential model

We model the mechanotransduction process based on the model published in Blum et al. 2020 and Simha & Ting 2024. We first compute the bag 1 and bag 2/chain components of the receptor potential. The bag 1 component of the receptor potential was computed as the weighted sum of the force and half-wave rectified time derivative of force, termed yank, from the bag 1 fiber. The yank was half-wave rectified to model the dynamic part of the receptor current as encoding only the positive rate of change of force. The bag 2/chain component of the receptor potential was computed as a weighted force from the bag 2/chain fiber. Finally, the total receptor potential of the Ia afferent was the sum of these two components.

### Integrate and fire model

The receptor potential is integrated until a threshold value is reached, at which point it generates a spike and undergoes a refractory period before it can generate a spike again. The time between each spike is estimated and the inverse of that time is reported as the model Ia firing rate.

### Simulations

All simulations were run as forward simulations where alpha motor activity impacted muscle fascicle length but did not drive it entirely, because muscle-tendon unit length was always imposed. Model parameters (Table 1) were manually tuned to fit a dataset from a published study where the Ia firing rate measured when sinusoidal ankle joint movements were imposed on an anaesthetized cat (39). All model parameters were then held constant in all simulations presented here, with only changes to the model input: MTU length, and alpha, gamma static, and gamma dynamic motor activations.

**Table 1:**
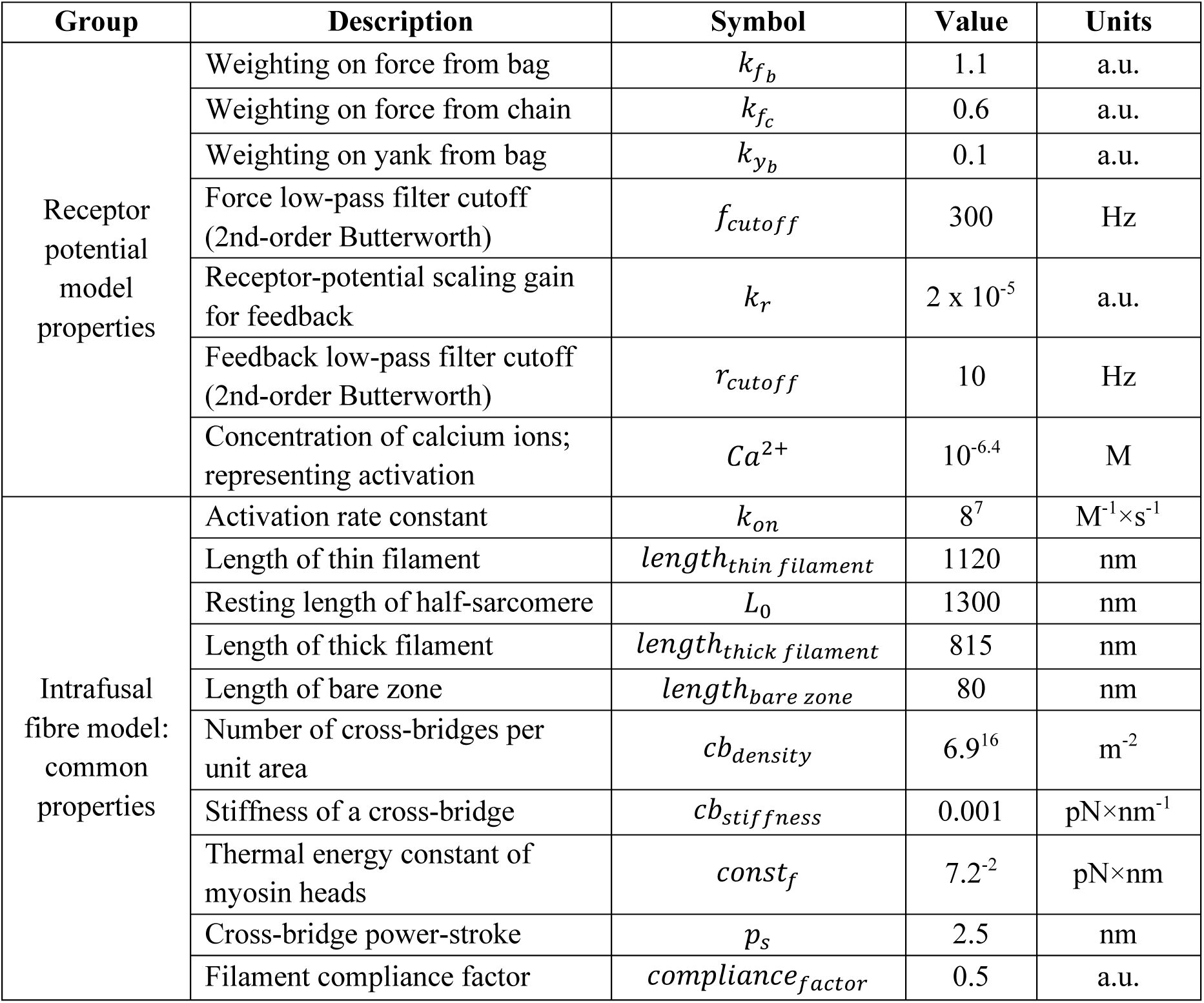
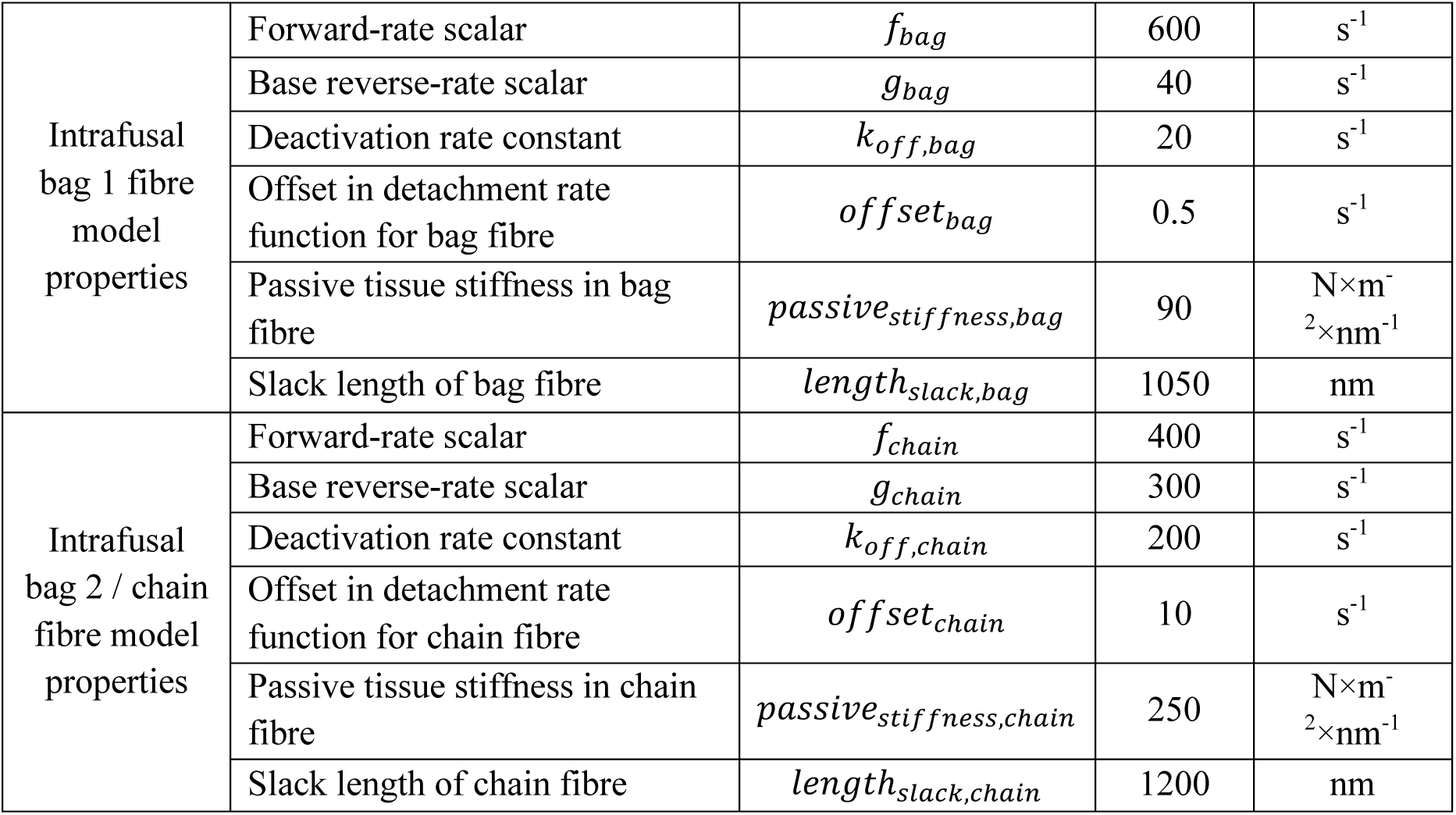
List of model parameters divided by receptor potential model and intrafusal fibre model. All values listed here were held constant through all simulations.

| Group | Description | Symbol | Value | Units |
| --- | --- | --- | --- | --- |
| Receptor potential model properties | Weighting on force from bag | $k_{fb}$ | 1.1 | a.u. |
| | Weighting on force from chain | $k_{fc}$ | 0.6 | a.u. |
| | Weighting on yank from bag | $k_{yb}$ | 0.1 | a.u. |
| | Force low-pass filter cutoff (2nd-order Butterworth) | $f_{cutoff}$ | 300 | Hz |
| | Receptor-potential scaling gain for feedback | $k_r$ | $2 \times 10^{-5}$ | a.u. |
| | Feedback low-pass filter cutoff (2nd-order Butterworth) | $r_{cutoff}$ | 10 | Hz |
| | Concentration of calcium ions; representing activation | $Ca^{2+}$ | $10^{-6.4}$ | M |
| Intrafusal fibre model: common properties | Activation rate constant | $k_{on}$ | $8^7$ | $M^{-1} \times s^{-1}$ |
| | Length of thin filament | $length_{thin\ filament}$ | 1120 | nm |
| | Resting length of half-sarcomere | $L_0$ | 1300 | nm |
| | Length of thick filament | $length_{thick\ filament}$ | 815 | nm |
| | Length of bare zone | $length_{bare\ zone}$ | 80 | nm |
| | Number of cross-bridges per unit area | $cb_{density}$ | $6.9^{16}$ | $m^{-2}$ |
| | Stiffness of a cross-bridge | $cb_{stiffness}$ | 0.001 | $pN \times nm^{-1}$ |
| | Thermal energy constant of myosin heads | $const_f$ | $7.2^{-2}$ | $pN \times nm$ |
| | Cross-bridge power-stroke | $p_s$ | 2.5 | nm |
| | Filament compliance factor | $compliance_{factor}$ | 0.5 | a.u. |
| Intrafusal bag 1 fibre model properties | Forward-rate scalar | $f_{bag}$ | 600 | $s^{-1}$ |
| | Base reverse-rate scalar | $g_{bag}$ | 40 | $s^{-1}$ |
| | Deactivation rate constant | $k_{off,bag}$ | 20 | $s^{-1}$ |
| | Offset in detachment rate function for bag fibre | $offset_{bag}$ | 0.5 | $s^{-1}$ |
| | Passive tissue stiffness in bag fibre | $passive_{stiffness,bag}$ | 90 | $N \times m^{-2} \times nm^{-1}$ |
| | Slack length of bag fibre | $length_{slack,bag}$ | 1050 | nm |
| Intrafusal bag 2 / chain fibre model properties | Forward-rate scalar | $f_{chain}$ | 400 | $s^{-1}$ |
| | Base reverse-rate scalar | $g_{chain}$ | 300 | $s^{-1}$ |
| | Deactivation rate constant | $k_{off,chain}$ | 200 | $s^{-1}$ |
| | Offset in detachment rate function for chain fibre | $offset_{chain}$ | 10 | $s^{-1}$ |
| | Passive tissue stiffness in chain fibre | $passive_{stiffness,chain}$ | 250 | $N \times m^{-2} \times nm^{-1}$ |
| | Slack length of chain fibre | $length_{slack,chain}$ | 1200 | nm |

#### Optimization

We used the MATLAB’s patternsearch function to identify the gamma motor activity that best matched the empirical data from Taylor et al., 2006 (18). We specifically needed to use a non-gradient based optimization algorithm because the gamma motor activation parameters generate a highly non-smooth cost landscape, leading gradient based searches to become stuck in local minima. We used eight optimization variables when the model inputs were entirely based on empirical data (Figure 1C,D): gamma dynamic activation magnitude when on, timing of gamma dynamic onset, timing of gamma dynamic offset, four points on a spline for gamma static, and a gamma static phase, relative to the MTU length. Optimization for the ideal sinusoidal input (Figure 1E) consisted of six optimization variables: gamma dynamic activation magnitude when on, timing of gamma dynamic onset, timing of gamma dynamic offset, gamma static activation vertical offset, gamma static maximum activation, and gamma static phase relative to alpha activity. The optimization was run in three stages to increase the probability of finding global minimum. In the first phase, the gamma dynamic timing variables were coarsely sampled and a patternsearch algorithm was run in parallel on the remaining variables, at each gamma dynamic timing pair. Phase 2 repeated the same process but using gamma dynamic timing pairs that were generated by finely sampling around the minimum cost variables found in phase 1. In phase 3 of the optimization, the gamma dynamic timing variables were fixed at the optimal values found in Phase 2, and seven random sets of initial values were generated for the remaining variables and run in parallel through patternsearch. The overall optimal values are reported. In the case of the ideal sinusoidal input (Figure 1E), muscle-tendon unit length was set to a 1.5Hz, 1% L0 amplitude sinusoid, while alpha activity was antiphase and ranged from 5 % to 65 % of maximum activation. We digitized the data from figures 3A, 4A, and 7B of Taylor 2006 (18) using WebPlotDigitizer (141). Both the digitized data as well as the model output were interpolated, and a mean-normalized root mean squared error used as the cost function for the optimization.

#### Efference copy simulations for gamma static

Gamma dynamic was kept at 0 % for all these simulations. Muscle-tendon unit length was a 1.5 Hz, 1% L0 amplitude sinusoid. Alpha motor activity was kept at zero for the simulation shown in Figure 2A and then set to be antiphase to the muscle-tendon unit and ranged from 35 % to 65 % of maximal activation for the other simulations shown in Figure 2. Gamma static activation was set to a tonic 25 % for the simulation in Figure 2A and 2B. Figure 2C shows two conditions of gamma static activity: gamma static is in-phase with alpha activity and either has an amplitude of 30 % or 13.5% of maximal activation, with a vertical offset of 25 %. For the results demonstrating the range of effects of gamma static on Ia firing rate, gamma static activity was swept through phase shifts of −50 % to +50 % relative to the gait cycle (muscle-tendon unit length), and through a range of amplitudes: being tonically active at 20% to a sinusoidal peak of 70% of maximal activation. Accordingly, two metrics are reported about the effect of gamma static modulation on Ia firing rate: the location of the peak firing rate relative to the gait cycle and the magnitude of the peak firing rate.

#### Feedforward versus feedback alpha motor activity

Gamma dynamic was kept at 0 % for all these simulations. Muscle-tendon unit length was a 1.5 Hz, 1% L0 amplitude sinusoid. The muscle spindle model was run one time step at a time such that the alpha activity that it received was determined as a combination of the feedforward alpha input as well as a scaled value of the muscle spindle Ia receptor potential from the previous time step. We used receptor potential rather than firing rate because the alpha motor activation to our model is provided as a continuous signal that is a percent of maximum activation, and therefore requires converting a firing rate value back into a continuous signal, which would introduce more processes and parameters. The simulation was run without feedback until 1.5 s to allow the forces to reach steady-state first. For all simulations in Figure 3, the feedforward alpha motor activity was set to range between 20 % and 76 % of maximal activation while gamma static activity ranged from 5 % to 95 %, and was in-phase with alpha motor activity for the first three simulations and antiphase for the fourth simulation. Total alpha motor activity was determined as a weighted sum of the feedforward and feedback alpha motor activity with feedforward having weights 1, 0, 0.5, 0.5 and feedback having weights 0, 1.5, 0.75, 0.75 for each simulation 1-4.

#### Perturbation experiments

To simulate the muscle spindle and consequent muscle fascicle response to perturbations, we generated a set of four simulations for every condition. 1) a baseline, unperturbed simulation 2) a simulation where the muscle-tendon unit was perturbed away from its sinusoidal motion for part of a cycle but no feedback was supplied to the alpha motor activity 3) a simulation where the muscle-tendon unit was perturbed away from its sinusoidal motion for part of a cycle and muscle spindle Ia receptor potential was supplied as feedback to alpha motor activity 4) a simulation where the muscle-tendon unit was perturbed away from its sinusoidal motion for part of a cycle and a proportional-derivative control signal designed to optimally correct the muscle fascicle deviation from baseline movement was supplied as feedback to alpha motor activity.

##### Perturbations

Two types of constant velocity perturbations were simulated. Both were length stretches applied to the muscle-tendon unit, in addition to the sinusoidal stretch-shorten cycles. The first was a slow stretch where the prescribed stretch magnitude was applied over a duration of 333 ms, with a ramp down over 50 ms to return to the baseline sinusoidal cycle. The second perturbation was a faster stretch where the prescribed stretch magnitude was applied over a duration of 50 ms. Figure 4A and 4B illustrate a 6 % L0 magnitude perturbation applied slowly (Figure 4A) and quickly (Figure 4B). The muscle-tendon unit sinusoidal cycles were always 1.5 Hz with 1 % L0 amplitude. Alpha motor activity was always antiphase with the muscle-tendon unit movement and ranged from 5 % to 65 % of maximal activation.

##### Proportional-derivative feedback control

The difference between the muscle fascicle length in the perturbed and matched unperturbed condition was estimated. A proportional-derivative signal was then generated as the weighted sum of the difference and its time-derivative. This process was performed at each time step of the simulation, and the proportional-derivative signal was added to the alpha motor activity at the following time step. The weights for the difference and its derivative, used in generating the proportional-derivative signal, were estimated as the values that minimized the root mean squared error between the unperturbed extrafusal fascicle length and when the muscle tendon unit was perturbed using a ramp of stretch magnitude 8 % L0, applied over a duration of 633 ms.

##### Muscle spindle feedback control

The baseline, steady-state muscle spindle Ia receptor potential obtained from the matched unperturbed condition was subtracted from the receptor potential generated during the perturbed condition, to obtain the muscle spindle feedback signal. This process allowed us to ignore introducing more parameters to determine the proportion of feedback and feedforward alpha motor activity in the baseline, unperturbed condition. The feedback signal was then filtered using a 10 Hz causal, low pass filter to account for activation dynamics and the closed-loop reflex bandwidth (142) and multiplied with a gain at each time step, and added as feedback to the alpha motor activity of the next time step. The gain on the muscle spindle Ia feedback was determined as the value that minimized the root mean squared error between the extrafusal muscle fascicle length with Ia feedback and the extrafusal muscle fascicle with proportional-derivative feedback when a 5Hz, 2.5 % L0 amplitude sinusoidal perturbation was applied to the muscle-tendon unit.

#### Perturbation experiments for gamma static

Gamma dynamic was held constant at 0 % for all the simulations in Figure 4. Figure 4A and 4B illustrate the feedback control effect in response to a perturbation magnitude of 6 % L0. Figures 4 C and D illustrate the effect across perturbation magnitudes ranging from 0 % to 10 % L0. Root mean squared deviation of the extrafusal muscle fascicle relative to the unperturbed condition was used as the metric to evaluate feedback control for the ramp perturbation since these are sustained perturbations. In contrast, the peak extrafusal fascicle error is reported for the ‘pulse’ perturbation where a rapid prevention of muscle stretch maybe favorable.

#### Perturbation experiments for gamma dynamic

These simulations illustrated in Figure 5 were equivalent to the ‘pulse’ perturbations illustrated in Figure 4, with the addition of gamma dynamic activity. Two conditions of gamma activity were simulated: one where gamma activity was absent, i.e., same as those in Figure 4, and one where gamma dynamic was phasically active, turning on at 4 % of the gait cycle and turning off at 18 %. The perturbation always occurred when gamma dynamic was active in the phasic condition, and was matched in location for the condition when gamma dynamic was absent. Figure 5 B and 5C illustrate the effect of gamma dynamic activity on Ia firing in response to perturbation as well as the peak extrafusal muscle fascicle deviation in response to the perturbation.

## Data Availability

All data used in the article were digitized from already published data. Simulation code will be made publicly available at https://github.com/Neuromechanics-Lab upon publication in a peer-reviewed journal.

## ACKNOWLEDGMENTS

We thank Friedl De Groote for helpful discussions and feedback at different stages of this study.

This work was supported by NIH NICHD R01 HD090642 to LHT, GSS, and TCC, and NIH NICHD R21 HD117352 to SNS and LHT.

